# Two CntT methyltransferases from *Actinosynnema mirum* and *Pseudomonas aeruginosa* reveal conserved opine-type metallophore methylation but divergent biological roles

**DOI:** 10.64898/2026.11.22.768471

**Authors:** Lola Bosc, Clémentine Laffont, Laurent Ouerdane, Thomas Beaurepere, Catherine Brutesco, Geneviève Ball, Pascal Arnoux, Romé Voulhoux

## Abstract

Opine-type metallophores have been discovered in mono- and diderm bacteria, where they allow zinc import under conditions of severe deficiency, a situation often found in natural environments. Whilst this family of metallophores is relatively well characterised, their modes of action in the organisms in which they are found remain poorly documented. In the present study, we identified and characterized two homologous SAM-dependent methyltransferases from the monoderm *Actinosynnema mirum* and the diderm *Pseudomonas aeruginosa* bacteria. We showed that the corresponding AmCntT (Amir_3913) and PaCntT (PA1921) enzymes methylate their cognate opine-metallophores, staphylopine and pseudopaline, respectively. Crystal structure of AmCntT established key structural features that are unique to this family of methyltransferase, and residues putatively involved in substrate stabilization were probed by site directed mutagenesis. Further investigation showed that, unlike the staphylopine from *A. mirum*, which is recovered in its methylated form in the extracellular medium, pseudopaline methylation occurred only under certain stress conditions and was only detected for an intracellular pool of pseudopaline, hence suggesting a role in its degradation in *P. aeruginosa*. Altogether, our data revealed a new common component of the *A. mirum* and *P. aeruginosa* opine-metallophore pathways respectively involved in staphylopine and pseudopaline methylation that appears to have distinct functional implications in each organism.

**IMPORTANCE:** Metallophores are small molecules with a very high affinity for metals. They are produced, secreted and recovered by bacteria in order to access metals under conditions of severe deficiency, a situation which is often encountered in the environment. In this study, we reveal the structure, function and role of two new methyltransferases involved in modifying the opine-type metallophores staphylopine and pseudopaline from *Actinosynnema mirum* and *Pseudomonas aeruginosa*. Contrary to the situation observed in *A. mirum*, where methylated staphylopine is detected in the extracellular medium, we showed that pseudopaline methylation was only detected intracellularly in *P. aeruginosa*. This suggests two different biological roles for these homologous enzymes. Our data illustrate the wide range of functions performed by metallophore methylation, including improved extracellular metal uptake, metallophore recycling, catabolism, and modulation of physicochemical properties at both the intracellular and microbial community levels.

## INTRODUCTION

Transition metals such as Fe, Zn, Cu, Mn, Ni and Co possess unique properties that make them essential for all living organisms [1]. Although these metals are ubiquitously present in the environment, their bioavailability for unicellular organisms such as bacteria, is often limited. As an initial response to metal scarcity, bacteria activate specific, high-affinity mechanisms for the uptake of remaining free metals [2]. To cope with more severe metal bioavailability, bacteria have developed powerful mechanisms relying on the production and secretion of molecules with high-metal affinity, called metallophores [3–6]. These means of metal recovery are notably important for pathogens, as the nutritional immunity settled by the host’s immune system diminishes the bioavailability of metals through the production of metal-chelating proteins [7, 8]. While transition metals are vital nutrients for bacteria, their presence in excess can also be highly toxic [2, 9]. The metal acquisition systems must therefore be tightly regulated to import metal only when required. This regulation is generally negative and mediated by a Fur- or Zur-type repressor whose inhibitory activity is proportional to the amount of intracellular metal. Thus, in the presence of metal, the repressor adopts an ‘active’ conformation able to recognize, bind and thereby block a specific DNA sequence – the *fur* or *zur* box – located upstream of the genes of interest [10].

The opine-type metallophore family has been described in several mono- and diderm bacterial species [11]. The first to be identified was staphylopine from *Staphylococcus aureus*, [12], and then pseudopaline from *Pseudomonas aeruginosa* [13, 14]. While staphylopine has been shown to have a broad range metal specificity in *S. aureus*, its function had clearly been linked to zinc homeostasis [15], despite the fact that its *cnt* operon is negatively regulated by both iron and zinc [16]. Concerning pseudopaline, it appears to be involved in the import of zinc in *P. aeruginosa*, playing a significant role in the acquisition of this metal under chelating conditions and during infection. [13, 14, 17, 18]. Additional opine-metallophores have later been identified in pathogenic bacteria from the *Yersinia* gender [19] and predicted in the soil bacteria *Paenibacillus mucilaginosus* and *Actinosynnema mirum* [11].

Metal acquisition by opine-type metallophores follows four sequential steps starting by the synthesis of the metallophore in the cytoplasm by CntK, CntL and CntM from S-adenosyl-methionine (SAM), histidine (either L or D isomers) and α-ketoglutarate (α-KG) or pyruvate; the histidine racemase CntK being restricted to opine-metallophores carrying D-histidine [20]. In the monoderm *S. aureus*, the neo-synthesized apo-staphylopine is secreted in the medium by the MFS transporter CntE before being recognized in its holo-form by the membrane-anchored substrate-binding protein CntA and imported in the cytoplasm by the ABC transporter CntBCDF [12]. In the diderm bacteria *P. aeruginosa*, the neo-synthesized pseudopaline is first exported in the periplasm by the drug metabolite transporter CntI before being taken in charge by the tripartite efflux pump MexAB-OprM which allows its final release in the extracellular medium. Once charged with zinc, the extracellular holo-pseudopaline is recognized and taken-up by the outer membrane TonB-dependent transporter CntO [13, 18, 21]. However, the mechanism by which the metal is subsequently extracted from this family of metallophores remains largely unknown. While Fe^3+^ is often released from siderophores *via* its reduction in Fe^2+^ [13, 18, 21, 22], Zn^2+^ must be released following a modification or degradation of the metallophore itself, as it does not possess a redox potential allowing its reduction. This stage of metal release is usually followed by either a recycling step of the metallophore or its degradation [23–26].

While all *cnt-*like operons possess genes encoding for synthesis, export and recovery of the metallophore, some of them contain additional genes encoding enzymes such as methyltransferases [11, 15, 27]. These enzymes catalyse the transfer of a methyl (-CH₃) group to diverse substrates, inducing changes in their physicochemical properties [28]. Different classes of methyltransferases can be found, but the most common is the SAM-dependent methyltransferases, which use a SAM molecule as methyl donor. Action of a methyltransferase in the biosynthesis of a siderophore is described for pyochelin, although the methyltransferase is embedded in the non-ribosomal peptide synthetase PchF [29]. The presence of predicted methyltransferase enzymes in several *cnt* operons suggests that they could play some roles in the opine-metallophore pathways, although this has never been investigated.

Here, we explore the roles, structures and functions of two methyltransferases (called CntT) associated with opine-type metallophores. In *Actinosynnema mirum*, the gene encoding this methyltransferase is the last gene in the staphylopine operon. We showed that methylated staphylopine could be detected in the extracellular medium. We showed that AmCntT could methylate staphylopine *in vitro*, and established its crystal structure in complex with S-adenosylhomocysteine. We further probed the role of conserved residues in the active site using site-directed mutagenesis. Unlike the situation described for *A. mirum*, pseudopaline methylation occurred only under certain stress conditions and was only detected for an intracellular fraction of pseudopaline, suggesting a role in pseudopaline degradation in *P. aeruginosa*.

## RESULTS

### Identification of methyltransferases in opine-type metallophores-producing organisms

Co-localisation of genes encoding methyltransferases with those involved in the biosynthesis of opine-type metallophores was early recognized [11, 27] although experimental approaches to decipher either their reactions or their eventual roles are still lacking. The *cnt* operon of *Actinosynnema mirum* encodes such a putative methyltransferase gene, located downstream the genes involved in metallophore biosynthesis, export, and recovery and called *cntT* (**Figure 1**). Composition of the *A. mirum cnt* operon resembles a combination of the one found in *Staphylococcus aureus* (sharing *cntKLM* and *cntE*) and in *Yersinia pseudotuberculosis* (sharing *cntPQR*), although the gene order is somehow rearranged (**Figure 1**). Perfecting this notion of gene fluidity, we noticed that the predicted CntT methyltransferase of *A. mirum* shares 41% sequence identity at the protein level with the gene product of *PA1921* from *P. aeruginosa,* located outside the *cnt* operon. Interestingly, *PA1921* was previously shown to respond to zinc limitation via Zur repression, alike the *cnt* operon [30], supporting a potential link between PA1921 methyltransferase and pseudopaline in *P. aeruginosa*. We therefore propose to rename these methyltransferases CntT and the following work is an attempt to decipher their role in the opine-type metallophore pathways in their corresponding bacteria.

**Figure 1:**
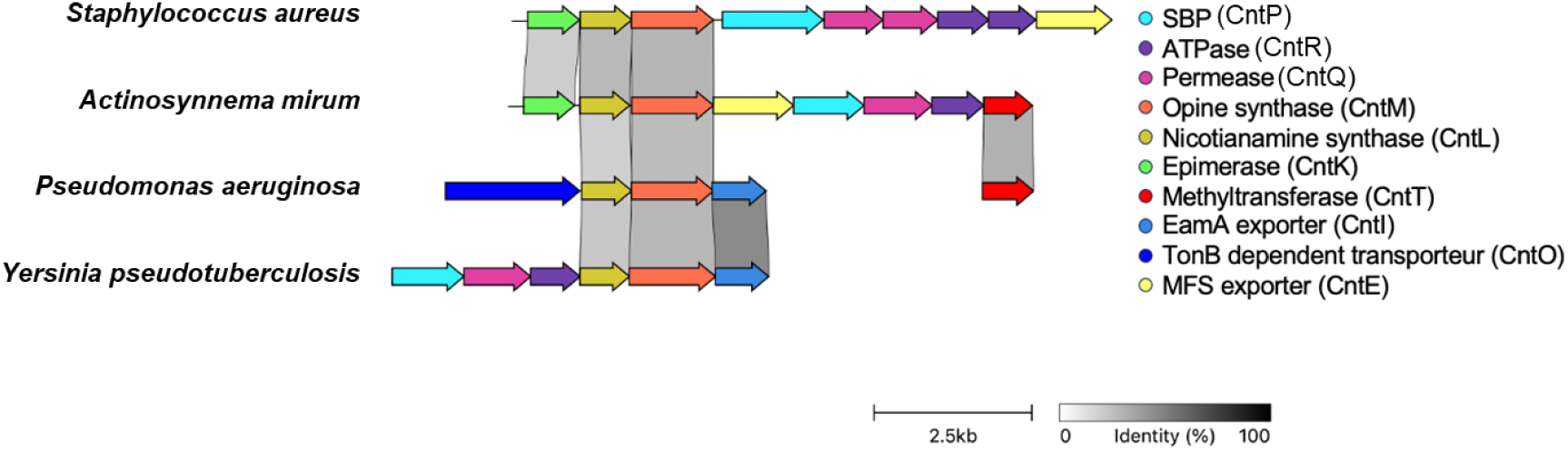
Comparative genetic organization of the *cnt* operons of *S. aureus*, *A. mirum*, *P. aeruginosa* and *Y. pseudotuberculosis.* The image is produced using the clinker program and the shades of grey depict protein sequence identity above 30%.

### Characterization of a SAM-dependent staphylopine methyltransferase in *A. mirum*

Based on the established rules to predict the identity of the produced opine-type metallophores [20], the presence/absence of a *cntK* gene - which determines whether D-His or L-His is incorporated - as well as the nature of the residue at position 150 of CntM (*S. aureus* numbering) - which determines whether pyruvate or α-KG is utilized - predicts that *A. mirum* is expected to produce staphylopine.

To confirm this, we grew *A. mirum* in metal chelating condition, collected the total cell and supernatant fractions and identified the opine-type metallophore using HILIC separation coupled to mass spectrometry (HILIC/ESI-MS). In both cellular and extracellular fractions, we were able to detect both staphylopine (Sp) and a methylated form of staphylopine (MeSp) (**Figure 2A**). *A. mirum* is therefore able to produce and secrete staphylopine and a methylated form of staphylopine.

**Figure 2:**
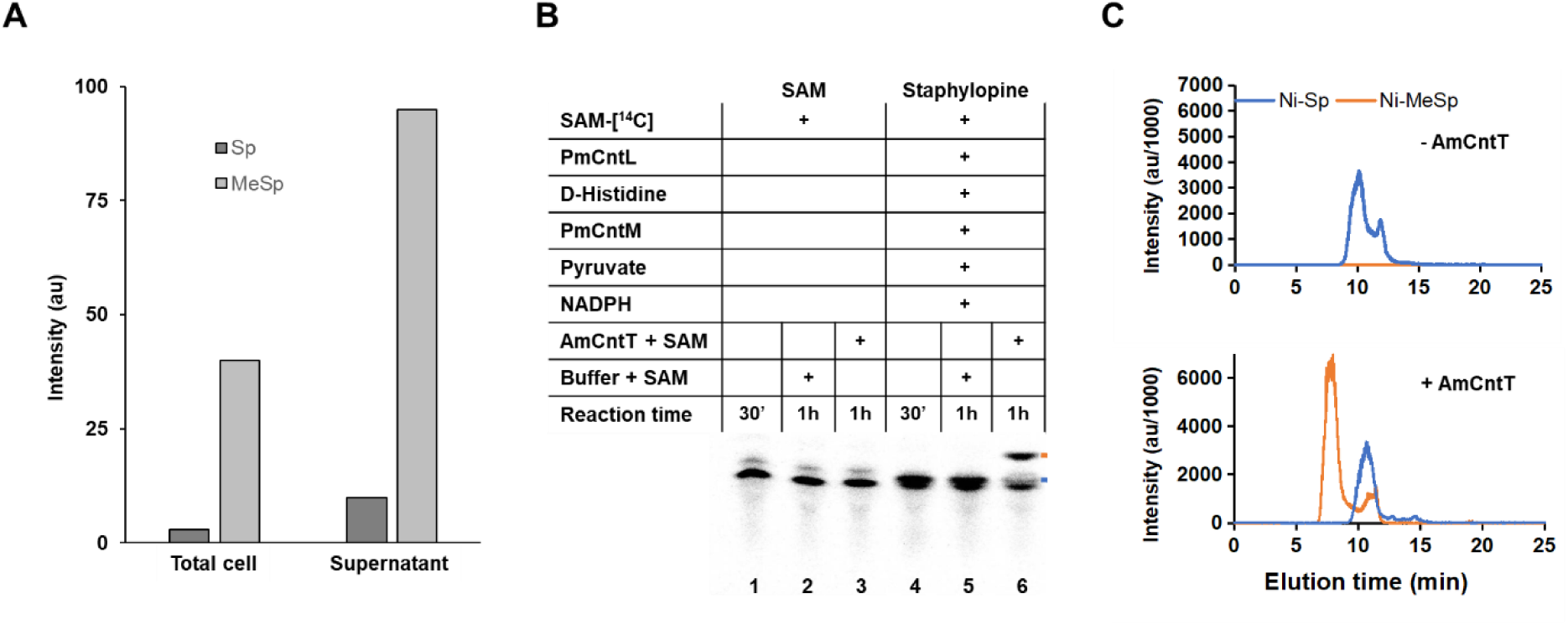
Methyl-staphylopine is produced by *A. mirum* using the methyltransferase AmCntT. (**A**) Detection of staphylopine (Sp) and methyl-staphylopine (MeSp) in the total cell fraction of a culture of *A. mirum* grown in TSB media supplement with 50 mM EDTA using HILIC/ESI-MS. (**B**) Production of radiolabelled staphylopine (blue mark) and methyl-staphylopine (orange mark) followed by TLC using 14C-SAM, enzyme mixes, substrates and cofactor. (**C**) Detection of staphylopine and methyl-staphylopine in the absence/presence of AmCntT by HILIC/ESI-MS.

To evaluate the role of the predicted AmCntT methyltransferase from *A. mirum* in the production of methylated staphylopine, we performed an *in vitro* reconstitution assays where radiolabelled staphylopine was produced using [¹⁴C]-SAM, D-histidine, pyruvate, NADPH and purified CntL and CntM from *P. mucilaginosus*, which were previously shown to efficiently produce staphylopine [20]. This staphylopine biosynthesis step was then followed by an additional incubation step with or without purified AmCntT and unlabelled SAM in order to assess methyltransferase activity (**Figure 2B**). The reaction mixtures were then analysed by thin-layer chromatography (TLC), as previously used to investigate opine-like metallophores [12, 13, 19]. Our results confirmed staphylopine production after 30 min of incubation, as indicated by a band pattern distinct from that of the control containing only [¹⁴C]-SAM (**Figure 2B blue mark, lane 4 versus lane 1**). In the presence of AmCntT and after incubation, a new band appears, which migrates further in the TLC (**Figure 2B lane 6, orange mark**). We attributed this band to methylated staphylopine, as no additional band appeared in the control reactions containing only SAM, with or without AmCntT (**Figure 2B lanes 2 and 3**), or in the reaction containing staphylopine without AmCntT (**Figure 2B lane 5**). To further confirm these findings, *in vitro* reactions containing staphylopine, with or without AmCntT, were analysed by HILIC/ESI-MS with prior spiking with nickel (**Figure 2C**). In the absence of AmCntT, only staphylopine was detected, whereas in its presence, both staphylopine and methylated staphylopine were detected. Together, these results demonstrate that, in *A. mirum,* AmCntT acts as a methyltransferase that uses staphylopine and SAM as substrates to produce and secrete methylated staphylopine.

### The three-dimensional structure of AmCntT reveals a new type of methyltransferase

Having established the activity of AmCntT, we next sought to decipher its three-dimensional structure in order to better characterize its structure-function relationship. We found that in solution, the purified protein elutes in size exclusion chromatography as a complex of approximately 150 kDa, consistent with either a pentamer or a tetramer. We were then able to obtain multiple crystals, among which one diffracted X-rays to 1.7 Å resolution. At the time, we solved its structure by the SAD method and were able to build the complete structure, encompassing residue N6 to E260 (for all monomers, except for monomer D, for which a loop between P216 and W222 is disordered) and one SAH molecule (the demethylated form of SAM) per monomer. The protein appears as a tetramer in the asymmetric unit, coherent with the size exclusion chromatographic profile (**Figure 3A, Figure S1**). Each subunit folds as a regular Rossman fold domain, with two additional features: a crown made of three α-helices (α6-α8), which contour the top of the domain where SAH is accessible, and three secondary structure elements (α9-β6-β7), which provide most of the residues involved in the tetrameric assembly of AmCntT (**Figure 3B**). The active site corresponds to the position of the Sδ atom of SAH (*ie* the position where the methyl group of SAM is transferred to its substrate). In AmCntT, this position is accessible to the solvent, and the electrostatic potential surface of this area suggests a rather deep binding pocket that is relatively apolar and delimited on one side by positive charge, whereas the 3-helices crown is more electronegative (**Figure 3C & 3D**). Residues building the ligand binding site are more or less conserved in a sequence alignment of diverse homologues of AmCntT (**Figure 3E**). In an attempt to probe this ligand binding site of AmCntT, we selected seven of these residues and systematically evaluated the effect of their corresponding alanine mutation on their activity. The TLC assay was then used to evaluate their capacity to produce methylated staphylopine (**Figure 3F**). We found that most site-directed mutants (Q19A, H130A, W131A, W191A and W215A) were unable to produce methyl-staphylopine whereas, two mutants (T127A and R211A) still retained the ability to produce it.

**Figure 3:**
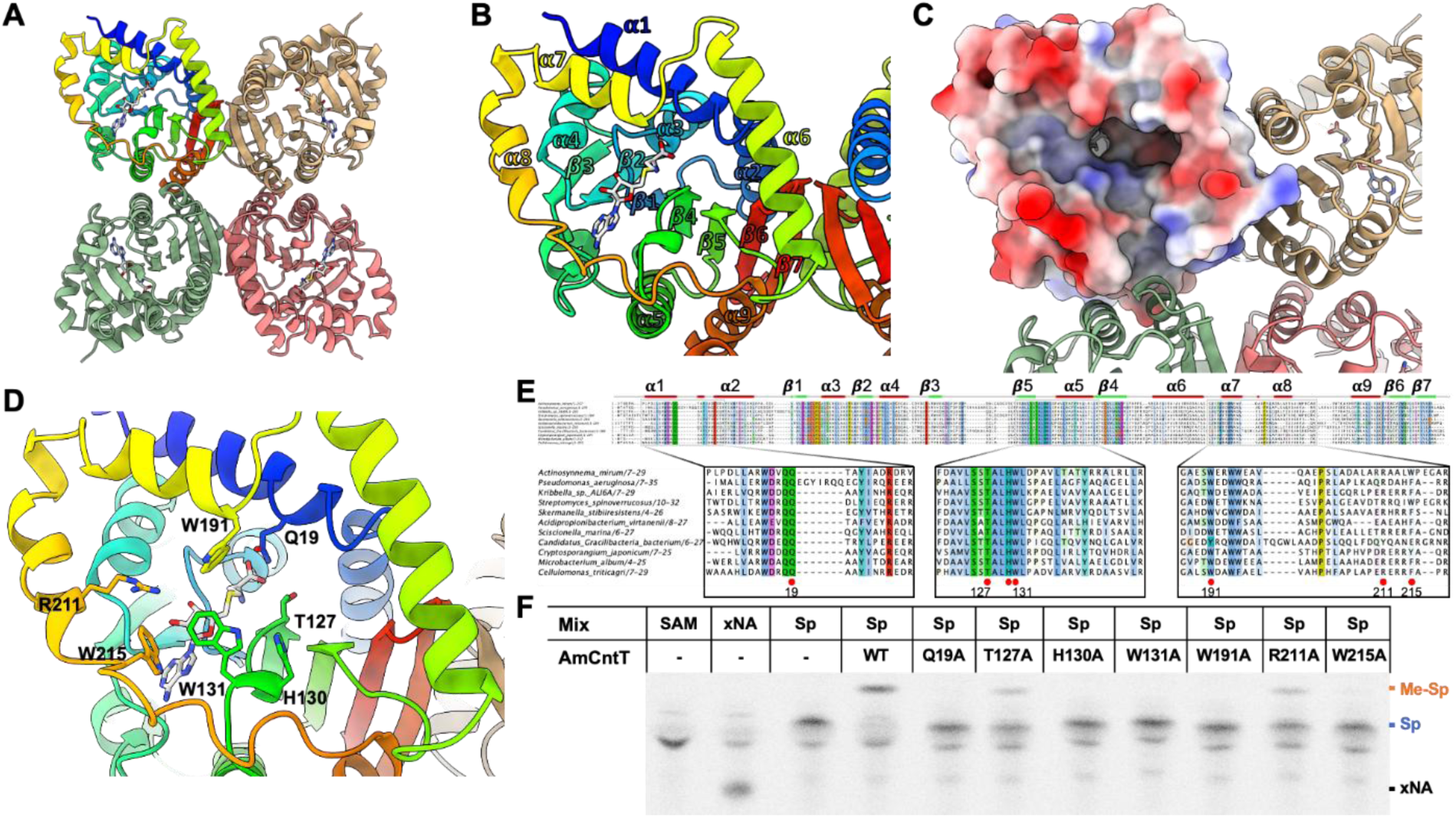
Three-dimensional structure of AmCntT and structure-function studies. **(A)** Tetrameric assembly of AmCntT, with one monomer coloured from blue (N-Terminus) to red (C-Terminus) and the other monomers in different colours. (**B**) Focus on one monomer and annotation of secondary structure elements highlights a three-helices crown (α6-α8) on top of the Rossman fold and three secondary structure elements (α9-β6-β7) that provide most of the residues involved in the tetrameric assembly of AmCntT. (**C**) Electrostatic potential of one monomer highlights a cavity in which SAH is accessible to the solvent and that likely accommodate its methyl acceptor substrate staphylopine. (**D**) The residues delineating this cavity are shown in stick. (**E**) Sequence alignment of diverse homologues of AmCntT, with zoom on parts of this alignment showing their sequence conservation. Residues selected for site-directed mutagenesis are highlighted by a red circle. (**F**) Production of radiolabelled staphylopine and methyl-staphylopine followed by TLC as in lane 6 of Figure 2, except that the production of methyl-staphylopine is probed by incubation with AmCntT WT protein or the indicated variants.

The AmCntT methyltransferase presents a conserved structure consisting of a Rossmann fold with a central cavity housing the active site composed of conserved residues. The closest structural homologue is a methyltransferase from *Methanosarcina mazei* (PDB code 3DTN) presenting 19.6% protein sequence identity, although a structural comparison show that the three-helices crown is not conserved and the quaternary structure is likely different (a monomer, possibly a dimer for 3DTN). Therefore, AmCntT represents a new type of methyltransferase.

### Identification of a CntT-family member in *Pseudomonas aeruginosa*

By analysing the genome of opine-type metallophore producing bacteria, an *A. mirum cntT* homolog was detected in *P. aeruginosa* strains. This gene called *PA1921* or *PA14_39660* respectively in the PAO1 and PA14 strains presents 59.3% nucleotide sequence identity with *cntT* of *A. mirum* while their corresponding proteins sequences are 41% identical and was therefore renamed PaCntT. Unlike in *A. mirum, cntT* of *P. aeruginosa* is not localized in the Zur-regulated *cntOLMI* operon that encodes for the synthesis, the export and the recovery of the opine-like pseudopaline metallophore by the bacterium (**Figure 1**). A Boltz structural model of PaCntT confirmed the homology with AmCntT with the model displaying a RMSD of 0.8 Å over 232 superimposed Cα atoms, including a conservation of the crown helices and of the last secondary structure elements involved in the quaternary assembly of AmCntT (**Figure 4A**). Furthermore, the identified active site residues of AmCntT (**Figure 3E**) appeared to be conserved in the central cavity of PaCntT (**Figure 4A**), suggesting a similar function for the two homolog proteins in the methylation of the metallophore produced by each organism. Although we could not determine the site of methylation mediated by PaCntT on pseudopaline (and neither of AmCntT on staphylopine) the predicted PaCntT-SAM-pseudopaline model indicated a proximity of the carboxylate of the L-histidine moiety of pseudopaline with the methyl group of SAM, suggesting that CntT belongs to the Class I of O-methyltransferase family [31, 32].

**Figure 4:**
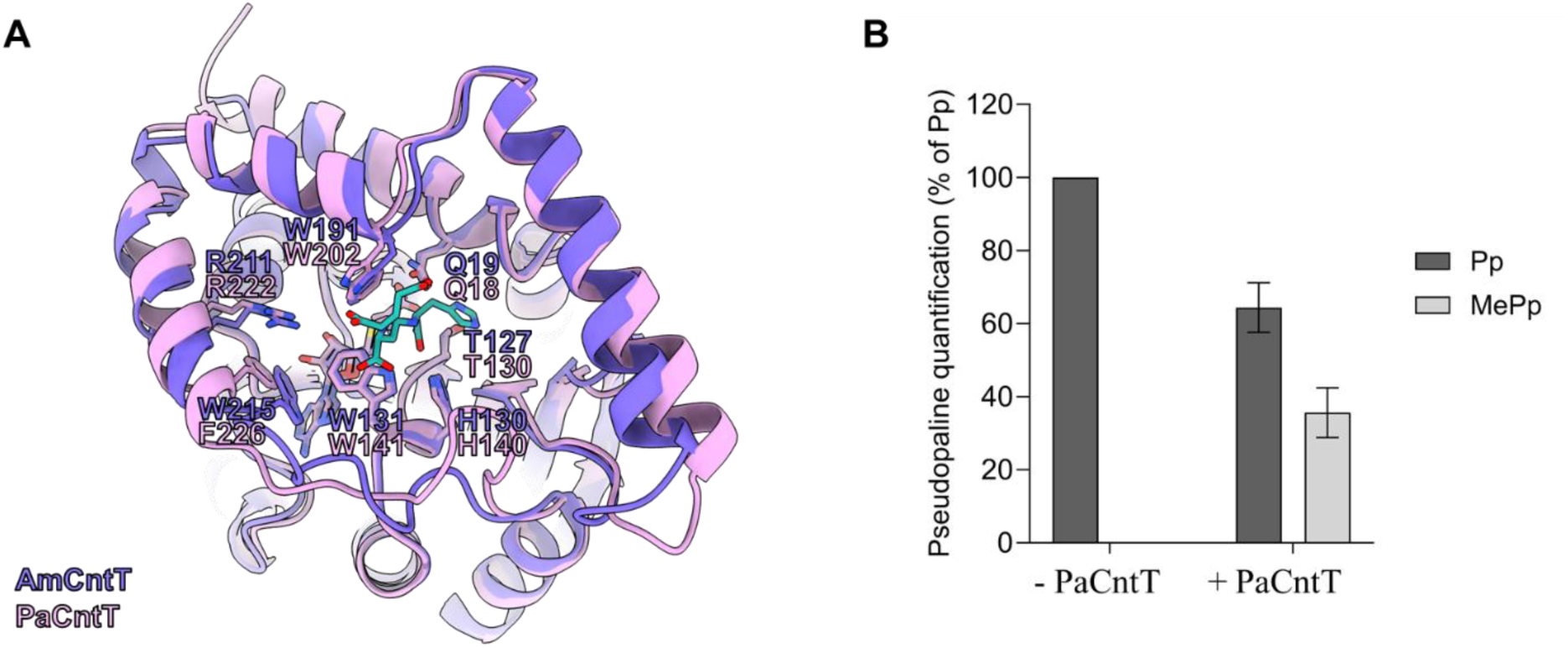
Identification and characterization of a CntT-family member in *P. aeruginosa*. **(A**) Overlay of the Boltz generated structural model of PaCntT with pseudopaline (salmon and green, respectively) with the AmCntT crystal structure (violet). Lateral chains of active site residues of AmCntT identified in Fig. 2 and their correspondence in PaCntT are represented. (**B**) HILIC-ESI/MS detection of pseudopaline (Pp) or methylated pseudopaline (MePp) following incubation of 100 μM synthetic pseudopaline with 5 μM purified PaCntT and 1 mM SAM. Error bars, mean ± standard deviation (SD) of at least three independent biological replicates.

We experimentally tested whether PaCntT could methylate the pseudopaline produced by *P. aeruginosa* by incubating the purified protein with the methyl-donor SAM and the *in vitro*-synthesized pseudopaline. Analysis of these samples by HILIC-ESI/MS revealed that in presence of PaCntT, 40% of the pseudopaline was methylated, confirming the enzymatic function of this protein and its ability to use pseudopaline as its substrate *in vitro* (**Figure 4B**). These results thus confirmed that alike AmCntT, PaCntT belongs to the opine-type metallophore methyltransferase family.

### Two different signals are necessary for *in vivo cntT* induction in *P. aeruginosa*

To assess whether PaCntT could be physiologically involved in opine-type metallophore methylation in *P. aeruginosa*, the conditions of its production were investigated. Metallophores of this family are generally subjected to metal-dependent regulation and only produced during metal starvation. As previously mentioned, Pederick and colleagues identified in *P. aeruginosa* a Zur-box upstream the *cntT* gene and found that it was upregulated in a Δ*znuA* mutant mimicking zinc scarce conditions [30]. To complete this observation, we assessed the production of PaCntT protein by immunoblot in a wild type or a Δ*zur P. aeruginosa* PA14 strain grown at 37°C in LB. In contrast to the pseudopaline outer membrane receptor CntO that reports the promoter activity of the *cnt* operon, PaCntT was surprisingly not detected in the Δ*zur* strain grown in rich medium LB, suggesting that the lifting of Zur repression alone is not sufficient to allow the expression of *cntT* (**Figure 5 upper left panel**). We therefore investigated whether the production of PaCntT was subjected to an additional layer of regulation other than zinc scarcity. Previous transcriptomic studies mentioned that *PA1921* (*cntT*) belongs to the RpoH (σ^32^) regulon [33]. This sigma factor is known to control the expression of genes involved in heat-shock response usually mimicked by incubation of the culture at 42°C. In accordance with this prediction, we found that the production of PaCntT in the Δ*zur* strain grown in LB medium was conditioned by a 42°C incubation, in contrast to CntO whose production was not temperature-dependent (**Figure 5 upper right panel**). It is worth noting that PaCntT is neither produced at 42°C in presence of the Zur repressor (in the WT strain) nor in absence of the repressor (in the Δ*zur* strain) without temperature shock, thus demonstrating that both signals are necessary to induce PaCntT production. It is also interesting to note that PaCntT is already detectable in the wild type strain grown in Minimal Chelated Medium (MCM), a metal-starved condition known to induce pseudopaline production [13], at 37°C, although its production is boosted at 42°C (**Figure 5 bottom panels**). This observation confirms that the first signal necessary for *cntT* expression in *P. aeruginosa* - zinc scarcity - is already effective in MCM compared to LB, therefore suggesting that heat-shock is an additional level of regulation of *cntT* in this bacterium. However, the observation that PaCntT is already produced in MCM without heat shock suggests that the physiological signal triggering its expression remains to be defined and that 42°C incubation might not be the only condition for the second layer of regulation. Altogether, these results show that PaCntT production requires zinc scarce conditions to relieve Zur repression, as well as an additional temperature-dependent induction or repression relief signal. This suggests a role in the global metal starvation response pathway, and the necessity of an additional signal for full gene expression indicates that PaCntT plays a role only under specific conditions that remain to be defined.

**Figure 5:**
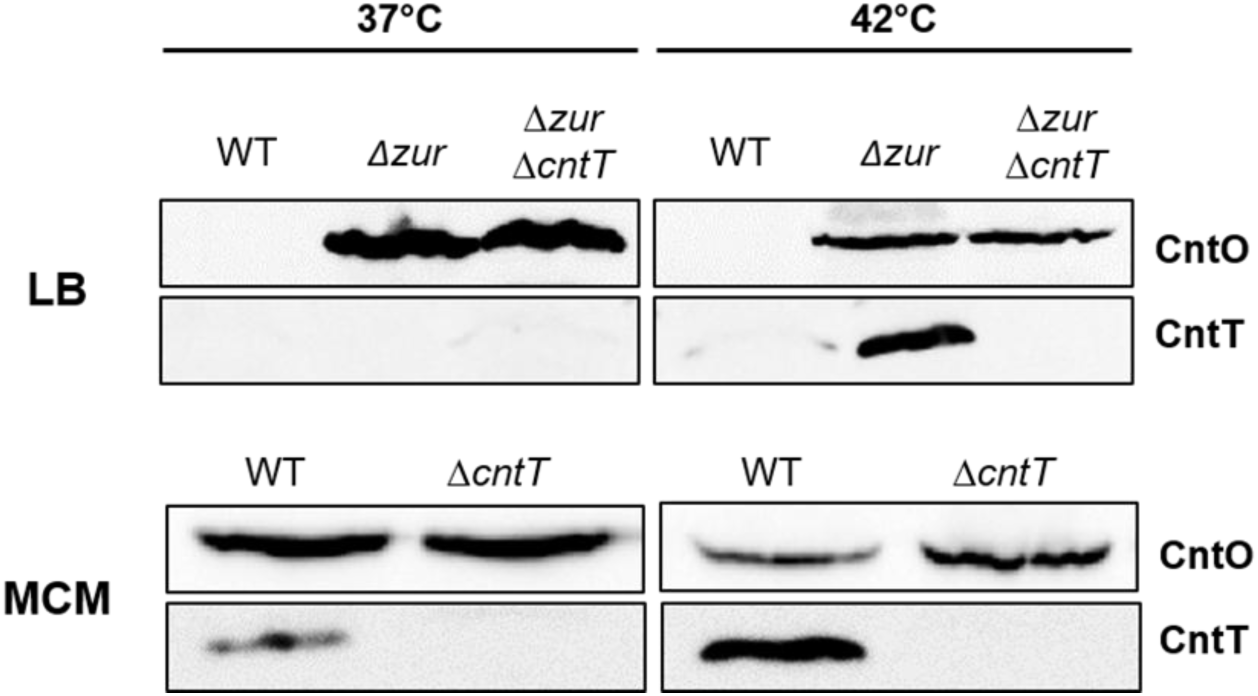
PaCntT production requires two signals. CntO and CntT protein detection by immunoblot in PA14 wild type (WT), *zur::tn* (Δ*zur*) and *zur::tn* Δ*cntT* (Δ*zur*Δ*cntT*) strains grown in LB or MCM medium at 37 or 42°C. Images presented are representative of at least two replicates.

### PaCntT methylates the pseudopaline accumulated in the cytoplasm, leading to its degradation

As PaCntT is able to methylate pseudopaline *in vitro* we hypothesized that, similarly to AmCntT, this methyltransferase could exert its function on the neo-synthesized pseudopaline prior to its secretion. To address this hypothesis, we analysed the supernatants of *P. aeruginosa* cultivated in MCM, a zinc-scarce condition known to be sufficient to induce the lifting of Zur repression and which allows a slight production of PaCntT (**Figure 5, lower left panel**). However, in order to warrant the production of PaCntT, considering that incubation at 42°C in MCM is a too harsh condition to allow sufficient amount of biomass for unbiased downstream analysis, we mimicked the increased production of PaCntT at this temperature by overproducing the protein from a plasmid harboring the *cntT* gene of *P. aeruginosa* under the control of an arabinose inducible promoter. Although pseudopaline is correctly produced and secreted, we were unable to detect any extracellular methylated pseudopaline, even under overproduction of the PaCntT methyltransferase (**Figure 6A**). Since PaCntT does not methylate the secreted pseudopaline, we investigated whether it could methylate the metallophore during the second stage of its cycle following its recovery for metal acquisition. Such hypothesis is supported by the intracellular detection of methylated pseudopaline in a non-producing (Δ*cntL*) and non-secreting (Δ*cntI*) PA14 strain incubated with exogenous pseudopaline (**Figure 6B**). The cytoplasmic localization of PaCntT (**Figure 6C**) further indicates that the observed pseudopaline methylation by PaCntT (**Figure 6B**) is performed in this compartment. These observations revealed that, in contrast to AmCntT in *A. mirum*, PaCntT of *P. aeruginosa* does not methylate the pseudopaline prior to its secretion it in the milieu, but rather after its way back to the cytoplasm. Our data moreover support the transient nature of the cytoplasmic stages of pseudopaline, as the neo-synthesized one, to which cytoplasmic PaCntT does not have access, might be quickly secreted and the imported pseudopaline-metal complex might be rapidly degraded and/or recycled.

**Figure 6:**
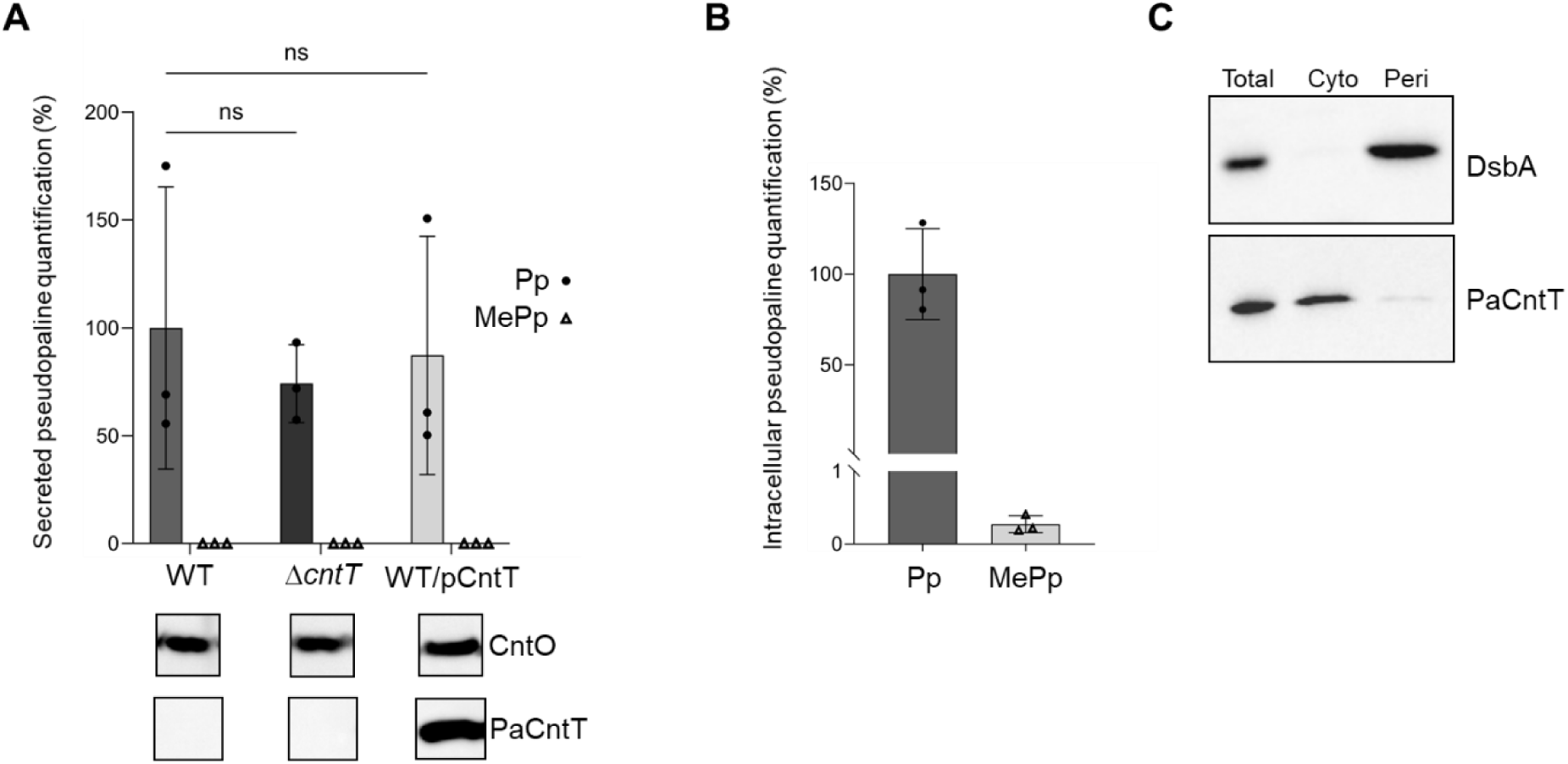
PaCntT selectively methylates the imported pseudopaline in the cytoplasm. (**A**) HILIC-ESI/MS quantification of pseudopaline (Pp) and methylated pseudopaline (MePp) in MCM culture supernatants of PA14 wild type strain containing the empty vector pJN105 (WT), Δ*cntT* pJN105 (Δ*cntT*) or wild type containing PaCntT-overexpressing plasmid pJN_CntT (WT/pCntT). Data are displayed as percentage of total pseudopaline in WT strain. Error bars, mean ± standard deviation (SD) of at least three independent biological replicates. The CntO and PaCntT production was assessed by immunoblot in each sample. Not significant (ns) (**B**) HILIC-ESI/MS quantification of intracellular total pseudopaline in PA14Δ*cntL*Δ*cntI* after 10 h growth in MCM and incubation for 10 minutes with 5μM pseudopaline-nickel complex following the procedure described in Bosc et al [17]. Results are represented as percentage of total pseudopaline. (**C**) Cellular localization of PaCntT by immunoblot using an anti-PaCntT antibody. The periplasmic protein DsbA serves as a control of fractionation. Intracellular total (Total), cytoplasmic (Cyto) and Periplasmic (Peri) fractions of a PA14 strain overexpressing PaCntT were examined by the two antibodies. Immunoblot images presented are representative of at least three replicates.

In order to further investigate the role of PaCntT in the *in vivo* methylation of the cytoplasmic pseudopaline, we measured the levels of methylated pseudopaline under conditions of PaCntT overproduction in the non-secreting (Δ*cntI*) strain, which accumulates the non-secreted pseudopaline in its cytoplasm [13]. Data presented in **figure 7A and B** indicated that the small but significant amount of intracellular methylated pseudopaline (0.5 % of total pseudopaline) is increased by 6-fold upon PaCntT overproduction, thus confirming the *in vivo* cytoplasmic methylation of pseudopaline by PaCntT. Based on these findings and the previous observation that the cytoplasmic accumulation of the neo-synthesized pseudopaline in the PA14Δ*cntI* strain is toxic for the bacteria [13], we further investigated the effect of its cytoplasmic methylation by PaCntT. Interestingly, we noticed that PaCntT overproduction significantly reduced the intracellular accumulation of total pseudopaline (**Figure 7A, left panel**), a phenotype also associated with a lifting of pseudopaline toxicity (**Figure 7C**). In line with these observations, the overproduction, to the same level as the wild type, of alanine variants of the two H133 and W195 active site residues of PaCntT (**Figure 3**) in the Δ*cntI* strain fully restore the toxicity phenotype, thus confirming that the reported PaCntT-dependent toxicity clearance is due to pseudopaline methylation by this methyltransferase (**Figure 7D**). These results therefore demonstrate that pseudopaline is either degraded or at least inactivated upon methylation by PaCntT which also explains why so little methylated pseudopaline is detected in the cytoplasm (**Figure 7A**), even under conditions of PaCntT overproduction. Altogether, these data revealed that PaCntT is able to methylate pseudopaline in the intracellular fraction. They moreover strongly suggest that this opine-type metallophore methylation step in *P. aeruginosa* is intended to degrade it or, at least, to inactivate it.

**Figure 7:**
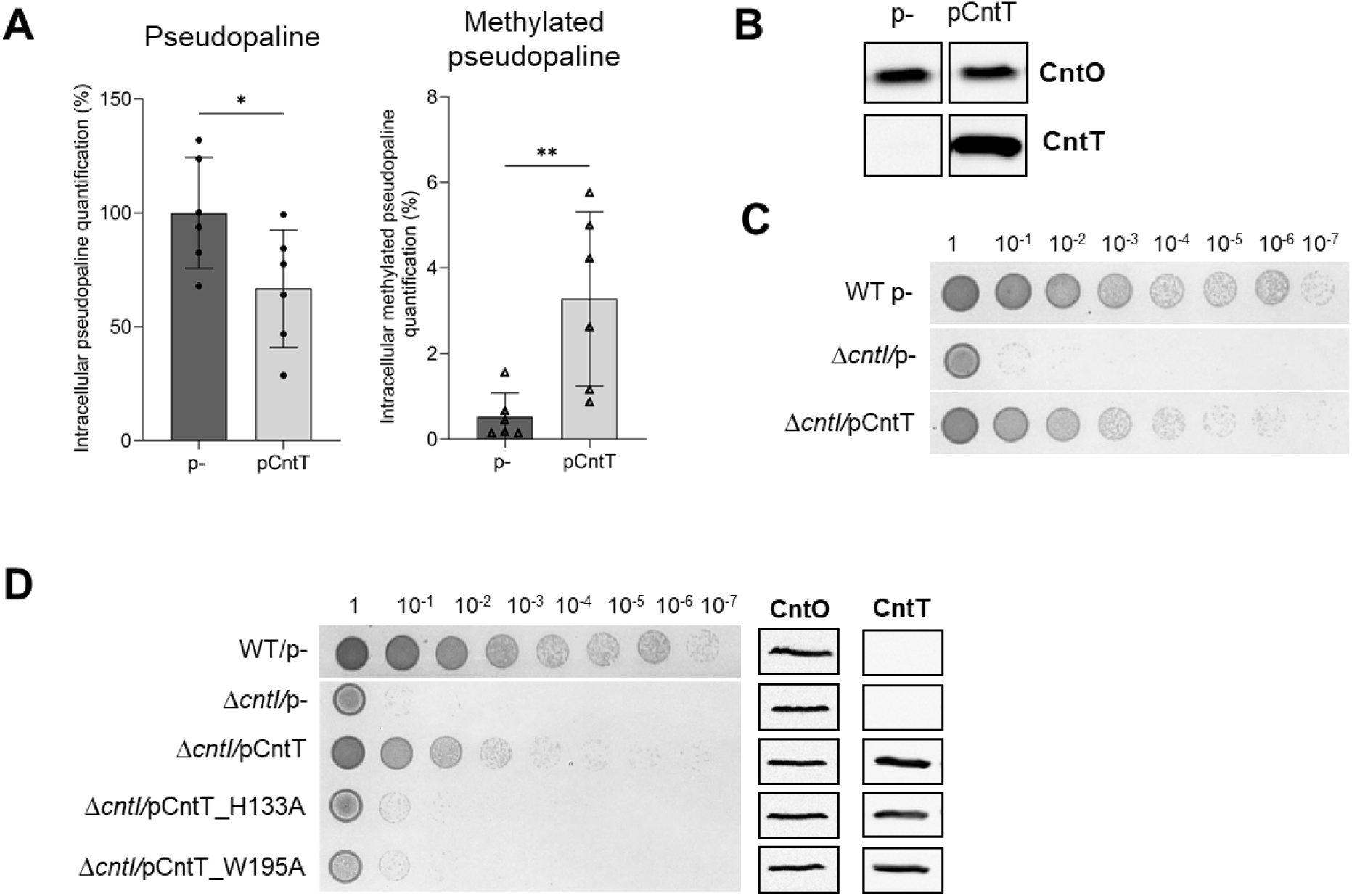
*in vivo* methylation of pseudopaline by PaCntT leading to its degradation. (**A**) HILIC-ESI/MS quantification of total pseudopaline or methylated pseudopaline in intracellular fractions of PA14 Δ*cntI* containing the empty vector pJN105 (p-) or pJN_CntT (pCntT) cultivated in MCM for 10 h. Data are displayed as percentage of total pseudopaline without CntT overproduction (p-strain). Error bars, mean ± standard deviation (SD) of at least three independent biological replicates. * and ** Correspond to p<0.05 and p<0.01 respectively (one-way ANOVA, Tukey post-hoc test). (**B**) CntO protein detection by immunoblot in samples from A. (**C**) Viability assay on plate of PA14 WT and Δ*cntI* strains containing the empty vector pJN105 (p-) or pJN_CntT (pCntT). (**D**) Viability assay on plate of PA14 WT and Δ*cntI* strains containing the empty vector pJN105 (p-), pJN_CntT (pCntT) or active site variants of CntT (pCntT_H133A, pCntT_W195A). Images presented in **B**, **C** and **D** are representative of at least three replicates.

### CntT is a versatile methyltransferase

The high sequence identity between AmCntT and PaCntT, combined with our observation that they methylate their respective opine-type metallophore prompted us to evaluate their substrate selectivity. Using a combined set of enzymes and substrates we were able to synthesized all the [^14^C]-labelled metallophores and, by co-incubation with AmCntT, to probe its substrate selectivity (**Figure 8A**). The migration on the TLC plate of all four metallophores were altered after co-incubation with AmCntT, indicating that the enzyme can use them all and therefore has a broad metallophore selectivity. The chemical modification is of the same nature, as all methylated forms migrated further on the TLC. Modelling of AmCntT with SAM and the four metallophores using Boltz indicates that they all fit in the same substrate binding site identified above, although in a different position (**Figure S2**). The same modelling was performed using PaCntT with the same conclusion, indicating that the metallophores could be used by the enzyme without a clear indication of position where the methyl group of SAM is transferred.

**Figure 8:**
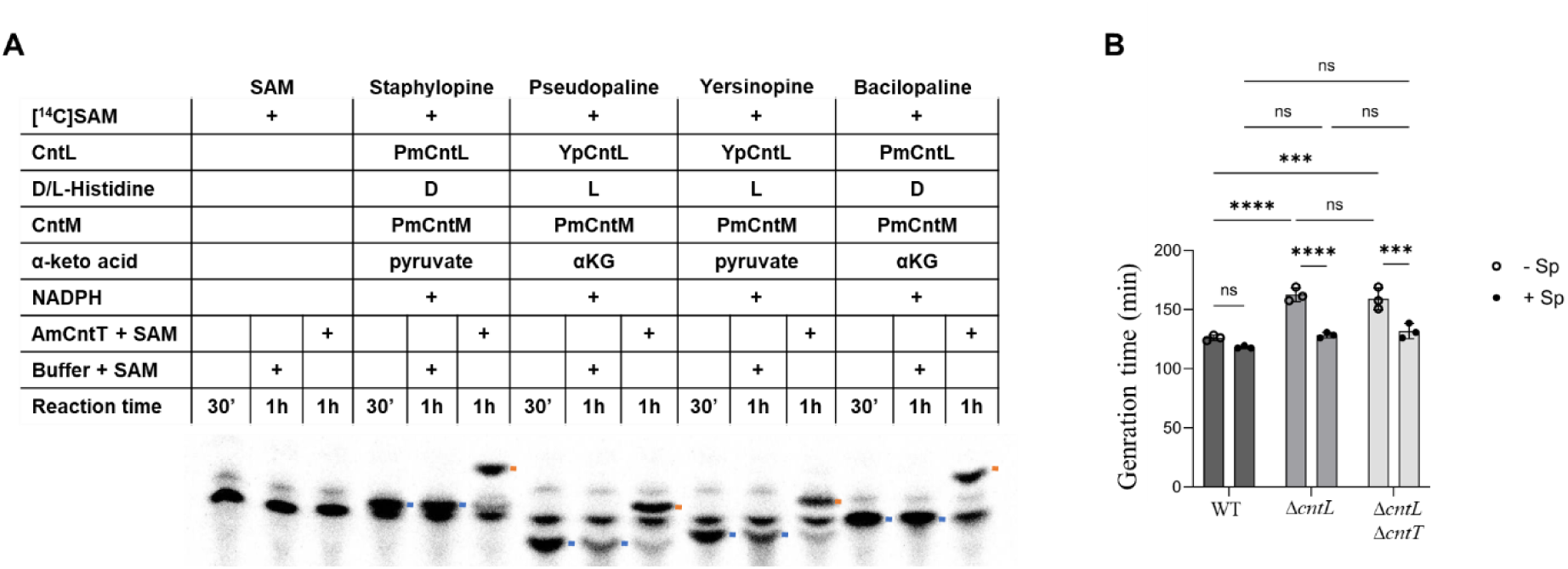
Broad specificity of AmCntT and cross-feeding of *P. aeruginosa* with staphylopine. (**A**) TLC separation of reaction products after incubation of [14C]-SAM with purified CntL and CntM enzymes from various organisms, α-keto acid, NADPH cofactor and L or D histidine. Co-incubation of both CntL and CntM enzymes with their substrates lead to the production of the corresponding opine-type metallophore, staphylopine, pseudopaline, yersinopine or bacillopaline, indicated by a blue dash. In presence of purified AmCntT and its methyl donor (SAM), an additional product, most probably corresponding to the methylated metallophore, is detected (red dash). (B) Generation time of PA14 WT, Δ*cntL* or Δ*cntL*Δ*cntT* strains cultivated in MCM supplemented or not with 5μM staphylopine. Error bars, mean ± standard deviation (SD) of at least three independent biological replicates. Not significant (ns); **** and *** Correspond to p<0.0001 and p<0.001 (one-way ANOVA, Tukey post-hoc test).

In order to test the cross-feeding of metallophores *in vivo*, we evaluated the putative impact of staphylopine supplementation on the growth of *P. aeruginosa*, for which a phenotype for the absence of pseudopaline corresponds to a longer generation time (**Figure 8B**) [17]. Although staphylopine supplementation had no effect on the generation time of the WT, it decreased it to a WT level in the PA14Δ*cntL* mutant, suggesting that *P. aeruginosa* benefited from staphylopine supplementation. However, since this exogenous staphylopine restoration phenotype is still visible in the PA14Δ*cntL*Δ*cntT* mutant, we conclude that PaCntT is not involved in the ability of *P. aeruginosa* to use staphylopine.

## DISCUSSION

In the present study, we identified a new family of SAM-dependent methyltransferases, called CntT, and found that they are involved in the methylation of opine-type metallophores. We investigated two CntT family members, AmCntT and PaCntT, and described their role in the *in vitro* and *in vivo* methylation of staphylopine and pseudopaline, respectively produced by the monoderm bacterium *Actinosynnema mirum* and the pathogenic diderm bacterium *Pseudomonas aeruginosa*. Additional experiments however revealed distinct role for the two CntTs in their respective organisms.

As methylated staphylopine is secreted by *A. mirum*, we hypothesized that this modification would play a role in extracellular metal acquisition. A main challenge for this bacterium forming multicellular aerial structures might be the transport of metals within aerial mycelium. Despite a current lack of evidence for the role of methyl-staphylopine in these structures, the transport of metals on long distances function of nicotianamine, an homologous metallophore produced by plants [34]. A similar role could thus be envisioned for methyl-staphylopine in *A. mirum*. Production of a methylated form of staphylopine could be an adaptation strategy for its diffusion in the aerial *vs* liquid environments in which the bacterium could grow. However, difficulties in genetically engineer *A. mirum* did not allow us to investigate in more detail the role of AmCntT methyltransferase in this bacterium.

In *P. aeruginosa,* while we confirmed *in vitro* that PaCntT is able to methylate pseudopaline, the supernatant of *P. aeruginosa* does not contain methylated pseudopaline even under PaCntT overproduction, ruling out a role of PaCntT in conferring a property to the extracellular secreted metallophore, alike it is proposed for *A. mirum*. This observation moreover revealed that before secretion, neo-synthesized pseudopaline is not freely accessible to the cytoplasmic PaCntT, thereby supporting the existence of a complex formed by synthesis enzymes and inner membrane transport pathway, allowing the immediate handling of newly synthesized metallophore by its internal membrane exporter, a mechanism similar to the one previously demonstrated for the siderophore pyoverdine [35, 36].

In *P. aeruginosa*, we however identified intracellular methylation of pseudopaline. We propose that this chemical modification probably leads to its degradation, which is supported by the decreased amount of cytoplasmic pseudopaline and the loss of its toxicity when accumulated in a *ΔcntI* mutant strain. Given these observations, it is only a short step to formulate the hypothesis that the role of PaCntT is to carry out the final step in the pseudopaline pathway, namely the release of the metal following its uptake by the bacterium. However, even if PaCntT does support this function, it must be restricted to the stress conditions where the protein is produced. Alternatively, PaCntT’s role could be to keep the cytoplasmic pseudopaline level below a threshold at which it could be toxic to the bacterium, for example, under heat stress conditions where the proteome and transport systems are disrupted [37] and in which PaCntT is produced. Preventing pseudopaline accumulation in these conditions might indeed help to prevents its toxicity.

In interplay between microorganisms mediated by a metallophore, *Staphylococcus aureus* was shown to be able to methylate the siderophore pyochelin produced by *P. aeruginosa* resulting in a reduced affinity for iron and a decrease toxicity toward *S. aureus,* which offers a dual advantage for *S. aureus* in presence of *P. aeruginosa* [38, 39]. This kind of metallophore modification could therefore have a strong impact on microbial communities. Considering that AmCntT is able to methylate all the known opine-type metallophores *in vitro*, one possibility is that PaCntT methyltransferase activity could also be versatile and target another opine-type metallophore, either to inhibit its activity for the producing organism, like it is the case for pyochelin, or to use it as a xenometallophore. *P. aeruginosa* is often found in co-infection, notably with the staphylopine producer *S. aureus*. By measuring the generation time of a PA14Δ*cntL* strain in metal scarce conditions, we demonstrated that the addition of exogenous staphylopine in the medium restored the growth of this strain. This effect was similar than the effect observed when providing exogenous pseudopaline to the Δ*cntL* mutant [17]. This observation therefore suggests that *P. aeruginosa* is able to alternatively use staphylopine for its growth in MCM. However, the additional deletion of *cntT* in a Δ*cntL* background does not impair this complementation, suggesting that PaCntT is not involved in this process under the used conditions. While not required for staphylopine utilization by *P. aeruginosa*, PaCntT could be involved in its methylation after import, in order to avoid its accumulation and potential toxic effects within the cell. It could also be involved in the modification of other opine-type metallophore in environments where metals constitute an essential resource, hence participating in the strong competition between microorganisms in these environments. More broadly, the mechanism highlighted in this study for PaCntT - metallophore methylation as a potential means of facilitating metal release following complex uptake - may extend beyond CntT-like methyltransferases. Indeed, analysis of the *A. mirum* genome revealed several additional methyltransferase genes (*Amir_4785*, *Amir_0328*, and *Amir_2704*) located in the vicinity of genes putatively involved in metal–complex import. Although the functions of these systems remain to be experimentally established, their genomic organization suggests that methyltransferase-mediated modification of imported metallophores may represent a more general strategy for metal recovery from xenometallophore complexes. Thus, metallophore methylation could represent a previously underappreciated mechanism contributing not only to metal acquisition, but also potentially to metallophore recycling, catabolism and the modulation of metallophore physicochemical properties at both intracellular and microbial community levels.

## MATERIALS AND METHODS

### Bacterial strains and growth conditions

Bacterial strains and plasmids used in this study are listed in Table 2.

All *P. aeruginosa* strains used in this study are derivatives of the parental PA14 strain. Unless otherwise specified, precultures of all *P. aeruginosa* strains were performed in minimal succinate (MS) medium (per liter: 6g K_2_HPO_4_, 3g KH_2_PO_4_, 1g (NH_4_)_2_SO_4_, 0.2g MgSO_4_, 4g succinic acid and 3.1g NaOH, pH 7.0) inoculated from MS + 1.5% agar and planktonic growths in minimal chelated medium MCM (MS medium supplemented with 100µM EDTA), both at 37°C with horizontal shaking in sterile disposable polycarbonate Erlenmeyer flasks in order to avoid any metal contamination and monitored by OD_600_ measurements. When needed, MS or MCM were supplemented with gentamycine 80 µg/mL and expression of *PacntT* from plasmid pJN_CntT was induced by adding arabinose 0.1%. External supplementation by chemical pseudopaline or staphylopine were respectively performed as described in [17] and [12].

*Actinosynnema mirum* DSM 43837 was used in this study. Cells were grown in liquid tryptic soy broth (TSB) diluted 10-fold relative to the supplier’s recommended concentration and supplemented with 50 µm EDTA. Cultures were incubated at 30°C for 48 h with horizontal shaking in sterile disposable polycarbonate Erlenmeyer flasks.

### Purification of AmCntT and variants, PaCntT, PmCntL, PmCntM and YpCntL

The genes encoding AmCntT and PaCntT were codon-optimized for *E. coli* expression, synthesized by Genewiz and cloned in pET-TEV between the NdeI and XhoI restriction sites, yielding plasmids pET-TEV-AmCntT and pET-TEV-PaCntT. Preculture of *E. coli* BL21/pET-TEV-PaCntT in LB kanamycine 25 µg/mL was used to inoculate a culture in terrific broth (BioBasic) kanamycine 2 5µg/mL. The expression of *PacntT* was induced by adding 0.1 mM IPTG before incubation overnight at 16°C with orbital shaking. The cultures were centrifuged (5000 g, 20 min, 4°C) and the pellets were resuspended in buffer A (Na_2_HPO_4_ 20 mM, NaCl 500 mM, imidazole 15 mM, pH 7.5) supplemented with DNAse 20µg/mL and antiprotease PMSF 1mM. Cells were lysed with French press and centrifuged a first time at low speed (4400 rpm, 20 min, 4°C). Supernatant was then ultracentrifuged (115,000 g, 45 min, 4°C) before 30 min incubation at 4°C in buffer A with Ni-NTA resin (Qiagen) previously washed with the same buffer. Proteins fixed to the resin were eluted with a gradient with increasing imidazole (buffer A supplemented with 250 mM imidazole). Fractions containing PaCntT were then transferred in buffer MES 50 mM NaCl 150 mM pH 6.5.

Preculture of *E. coli* BL21/pET-TEV-AmCntT and the variants investigated in this study (Q19A, T127A, H130A, W131A, W191A, R211A, and W215A) were used to inoculate cultures in terrific broth with kanamycine 50 µg/mL. The protein expression was induced by adding 0.1 mM IPTG before incubation at 16°C with orbital shaking. After overnight growth, the cultures were centrifuged (5000 g, 20 min, 4°C) and the pellets were resuspended in buffer A (Na_2_HPO_4_ 20 mM, NaCl 500 mM, imidazole 15 mM, pH 7.5) supplemented with DNAse (100 µg/L) and antiprotease (50 µl/L). Cells were lysed using a French press at 1000 psi (Aminco) and centrifuged (9000 g, 15 min, 4°C). Supernatants were ultracentrifuged (115000 g, 45 min, 4°C). The resulting soluble fractions were purified by batch using a nickel-charged affinity resin (Ni-NTA Agarose resin, ThermoFisher Scientific) previously washed with the same buffer A. Proteins fixed to the resin were eluted stepwise with buffer A supplemented with imidazole (15 mM to wash and 250 mM to elute). Purified proteins were finally transferred in an imidazole-free buffer (Hepes 20 mM, NaCl 300 mL, pH 7.5) using PD10 desalting columns. YpCntL, PmCntL and PmCntM, used to produced opine-type metallophores *in vitro*, were purified similarly with only variation in buffer A used to wash (potassium phosphate 50 mM, NaCl 300 mM, Imidazole 50 mM, pH 8 and Na_2_HPO_4_ 20 mM, NaCl 500 mM, imidazole 15 mM, pH 7.5), elute (potassium phosphate 50mM, NaCl 300 mM, Imidazole 250 mM, pH 8 and Na_2_HPO_4_ 20 mM, NaCl 500 mM, imidazole 500 mM, pH 7.5) and store (potassium phosphate 50 mM, sodium citrate 150 mM, pH 8, 10% glycerol and Hepes 20 mM, NaCl 300 mL, pH 8) proteins, as already described (Laffont, BiochemJ 2019; Laffont, Biometals 2026).

### *In vitro* methyltransferase assays

5µM of purified PaCntT was incubated for 30 min at 37°C with 5µM S-adenosyl-methionine (Sigma) and 100µM of synthetic pseudopaline (synthesis described in [40]) in 50mM ammonium acetate pH 7.0. Samples were conserved at -80°C before analysis by HILIC/ESI-MS as described later.

To study AmCntT activity *in vitro*, we proceeded in two steps. First, opine-type metallophores were produced using purified proteins and their respective substrates. Based on their substrate specificities, three enzymes from *Yersinia pseudotuberculosis* (Yp) or *Paenibacillus mucillaginosus* (Pm) were used to produce the four metallophores: YpCntL for reactions incorporating L-histidine, PmCntL for reactions incorporating D-histidine and PmCntM for all reactions, incorporating either pyruvate or α-ketoglutarate. The resulting combinations yielded staphylopine (D-histidine and pyruvate), pseudopaline (L-histidine and α-ketoglutarate), yersinopine (L-histidine and pyruvate) and bacillopaline (D-histidine and α-ketoglutarate). Reaction mixtures contained purified YpCntL or PmCntM (10 µM), purified PmCntM (10 µM), L- or D-histidine (10 µM), [^14^C]-SAM (2.5 µM), pyruvate or α-ketoglutarate (1 mM), NADPH (30 µM) in ammonium acetate buffer (100 mM, pH 7). Reactions were incubated at 28°C for 30 min. Second, mixtures were supplement with cold SAM (50 µM) and (or not) purified AmCntT (10 µM). Reactions were the incubated for additional 30 min at 28°C. The reactions were stopped by adding ethanol to a final concentration of 50% (v/v) and the products were then separated by a thin-layer chromatography. An aliquot of 10 ml of the reaction mixtures were spotted on HPTLC (High-Performance TLC) Silica Gel 60 Glass Plates (Merck KGaA), and the plates were developed with a phenol:n-butanol:formate:water (12 : 3 : 2 : 3 v/v) solvent system. HPTLC plates were dried and exposed to a [^14^C]-sensitive imaging plate for one day. Imaging plates were then scanned on a Typhoon FLA 7000 phosphorimager (GE Healthcare).

Alternatively, samples were prepared for analysis by HILIC/ESI-MS. In this case, staphylopine was first produced by incubating PmCntL (2.5 µM), PmCntM (2.5 µM), cold SAM (0.2 mM), D-histidine (0.2 mM), pyruvate (1mM) and NADPH (0.2 mM) in ammonium acetate buffer (100 mM, pH 7) for 30 min at 28°C. Reaction mixtures were then supplemented with cold SAM (0.2 mM) and (or not) purified AmCntT (5 µM), and incubated for additional 30 min at 28°C. The reactions were stopped by freezing and samples were conserved at -80°C until analysis.

### Pseudopaline and staphylopine identification and quantification by HILIC/ESI-MS

For *in vivo* pseudopaline quantification, *P. aeruginosa* cultures incubated for 10 h at 37°C under horizontal shaking in MCM were centrifuged 20 min at 4,500 rpm, 4°C. Supernatants were filtered at 0.2 µm and stored at -80°C before analysis. Pellets are washed three times with fresh MS medium and resuspended in ammonium acetate 20mM pH 7.0. Cells were lysed by sonication (VialTweeter, 3 cycles of 30s on/off during 5 min with 30s off between each cycle, C 70%, A 95%) and centrifuged at 16,000 g for 10 min, 4°C. Supernatants constituting intracellular fraction were stored at -80°C. These samples were analysed for pseudopaline quantification using HILIC/ESI-MS as previously described [13]. Pseudopaline levels were determined using the relative intensity of the m/z corresponding to the pseudopaline-Ni complex in samples saturated with nickel, in order to force nickel complex formation. For intracellular imported pseudopaline quantification, PA14 Δ*cntL*Δ*cntI* strain was cultivated as described previously. 5 µM of synthetic pseudopaline preincubated with NiCl_2_ (1:1) were added to the culture 10 min before centrifugation.

For measurements of staphylopine in intra- and extracellular fractions of *A. mirum,* cells grown for 48h at 30°C in 1/10 TSB supplemented with 50 µM EDTA were centrifuged (5000 g, 20 min, 4°C). Supernatants were filtered (0.22µm) to form the supernatant fractions, while cell pellets were washed in ammonium acetate buffer (20 mM). Cells were lysed using a French press at 1000 psi (Aminco) and centrifuged (9000 g, 20 min, 4°C). Resulting supernatants were filtered (0.22 µM) to form the total cell fractions. These samples, together with those containing *in vitro*-produced staphylopine with or without AmCntT were analysed as previously described using HILIC/ESI-MS. Relative intensity of the m/z corresponding to staphylopine-Ni complex (385.1) and methylated staphylopine-Ni complex (399.1) were measured.

### Structure determination of *Am*CntT

Similarly to WT AmCntT, selenomethionine-labelled AmCntT was produced by growing cells in SelenoMethionine Medium Base supplemented with SelenoMet Nutrient Mix (Molecular Dimensions), 40 mg selenomethionine, and 100 mg each of L-lysine hydrochloride, L-threonine and L-phenylalanine. After purification, proteins remained fused to a TEV tag. This fusion was cleaved by diluting the protein to 1 mg/mL in buffer A (Na_2_HPO_4_ 20 mM, NaCl 500 mM, imidazole 15 mM, pH 7.5) and incubating overnight with TEV protease at a mass ratio of 1:80. Following cleavage, the protein solution was passed through a nickel-affinity column, and the cleaved protein was collected in the flow-through by washing the column with buffer A. The collected fraction was then loaded onto a HiLoad 26/60 Superdex 200 size-exclusion chromatography column (GE Healthcare) equilibrated with buffer C (20 mM HEPES, 50 mM NaCl, pH 8.0).

Crystallisation conditions were screened using commercial crystallisation screens by sitting-drop vapour diffusion. Equal volumes (0.7 μL) of purified selenomethionine-labelled AmCntT (concentrated at 10mg/mL) and crystallisation solutions were mixed using a Freedom EVO liquid-handling robot (Tecan). The protein crystallised using 1.9 M ammonium sulfate, 0.1 M Tris and 20 mM xylitol, pH 8.5. Crystals were harvested, transferred to the crystallisation buffer supplemented with 20% glycerol and flash-cooled in liquid nitrogen. Diffraction data were collected at the SOLEIL synchrotron (Saint-Aubin, France, Table 1).

**Table 1.** Data collection and refinement statistics.

| **Data collection** |  |
|---|---|
| Wavelength | 0.9789 |
| Resolution range | 45 - 1.7 (1.761 - 1.7) |
| Space group | *P*2<sub>1</sub>2<sub>1</sub>2<sub>1</sub> |
| Unit cell | 77.1 86.6 159.2 90 90 90 |
| Total reflections | 234437 (22946) |
| Unique reflections | 117287 (11528) |
| Multiplicity | 2.0 (2.0) |
| Completeness (%) | 99.63 (99.12) |
| Mean I/sigma(I) | 23.39 (2.17) |
| R-merge | 0.02159 (0.3317) |
| CC1/2 | 0.999 (0.862) |
| **Refinement** |  |
| R-work | 0.1732 |
| R-free | 0.2043 |
| CC(work) | 0.961 |
| CC(free) | 0.965 |
| Number of non-hydrogen atoms | 8899 |
| Macromolecules | 7983 |
| Solvent | 916 |
| Protein residues | 1014 |
| RMS(bonds) | 0.009 |
| RMS(angles) | 1.11 |
| Average B-factor | 30.61 |
| Macromolecules | 29.70 |
| Solvent | 38.59 |

Statistics for the highest-resolution shell are shown in parentheses.

Selenium sites in the selenomethionine-labelled protein were identified using the SHELX suite [41]. The resulting electron-density maps were used for initial model building with Buccaneer [42]. Model inspection and rebuilding were performed using Coot [43], and refinement was carried out with phenix [44]. Structural visualisation and analysis were performed using ChimeraX [45]. Structure of CntT was deposited to the Protein Data Bank (pdb code: 35NQ).

### Viability assay on plate

Overnight MS precultures of PA14 strains were used to inoculate a culture of fresh MCM supplemented with gentamycine 80 µg/mL at OD600 of 0.1 and incubated under horizontal shaking at 37°C. The expression of *PacntT* was induced by adding arabinose 0.1% to the culture. After 9 hours, cultures were adjusted to OD600 of 1 and subjected to 10% serial dilutions in fresh MCM. 5µL of culture samples were spotted on LB 1.5% agar plates and incubated overnight at 37°C.

### Protein detection by immunoblot

For analysis of CntO and PaCntT production, protein samples from bacteria corresponding to 1 OD unit were resuspended in 50µL 1X SDS-PAGE loading Laemmli buffer containing β-mercaptoethanol, heated at 95°C for 10 min and 5µL was loaded and separated on 12% acrylamide SDS-PAGE. After separation, the proteins were transferred onto nitrocellulose membranes for western blot analysis. Transfers were performed with semi-dry technic using Power Blotter Station (Invitrogen). The membranes were washed briefly in Tris-buffered saline, 0.10% Tween 20 (TBS-T) and then blocked in TBS-T + 10% milk for 1 hour at RT. The membranes were incubated for 1h at RT in TBS-T + 10% milk containing the primary mouse or rabbit antibodies directed respectively against CntO, PaCntT or DsbA (laboratory collection), respectively at 1/1,000, 1/500 and 1/1,000 dilution. This was followed by 4 washes in TBS-T before an incubation of 45 min at RT in TBS-T + 10% milk containing the secondary antibody anti-mouse or anti-rabbit HRP conjugated (1/5,000). Finally, after five more washes with TBS-T, the peroxidase reaction was revealed by chemiluminescence (SuperSignal^TM^ West Dura Extended Duration Substrate, Thermo Scientific) and scanned with ChemiDoc (Bio-Rad). Cell fractionation of cytoplasmic and periplasmic compartments was realized as described by Ize *et al.* [46].

### P. aeruginosa genetics

Plasmids used in this study are listed in Table 2. Polymerase chain reactions (PCRs) were performed using the Q5 high-fidelity DNA polymerase (New England BioLabs) for routine cloning or Pfu Turbo (Agilent) for quick-change mutagenesis. Custom oligonucleotides, listed in Table 2, were synthesized by Integrated DNA Technologies. For *ΔPacntT* deletion mutant generation, two DNA fragments corresponding to upstream and downstream regions of *PacntT* gene were amplified from PA14 chromosome DNA with PCR primers LB-27/LB-28 and LB-29/LB-30 (Table 3). Upstream and downstream regions were ligated by overlapping PCR and cloned into linearized pKNG101 [47] by the SLIC method [48]. The resulting constructs were transformed into *E. coli* CC118λ*pir* and introduced into *P. aeruginosa* PA14 by triparental conjugation using the helper strain 1047/pRK2013. The strains in which the chromosomal integration event occurred were selected on *Pseudomonas* isolation agar plates supplemented with 2 mg/mL streptomycin. Excision of the plasmid, resulting in the deletion of the chromosomal target gene, was performed after selection on Luria-Bertani (LB) plates containing 6% sucrose. Sucrose resistant and streptomycin sensitive clones were confirmed to be deleted for *PacntT* gene by PCR analysis using LB-31/LB-32. The Δ*cntI* Δ*cntT* strain was constructed by knocking-out *PacntT* in the Δ*cntI* strain. *PacntT* overexpression was achieved by amplifying *PacntT* gene from PA14 chromosomic DNA using primers LB-60/LB-61 and cloning by SLIC in the vector pJN105. The construction of alanine variants was done by quick-change mutagenesis using pJN_CntT as template and the primers LB-95/LB-96 and LB-97/LB-98 listed Table 3. All constructs have been verified by DNA sequencing (Eurofins Genomics). The plasmids obtained were introduced in *P. aeruginosa* by triparental conjugation as previously described.

**Table 2:**
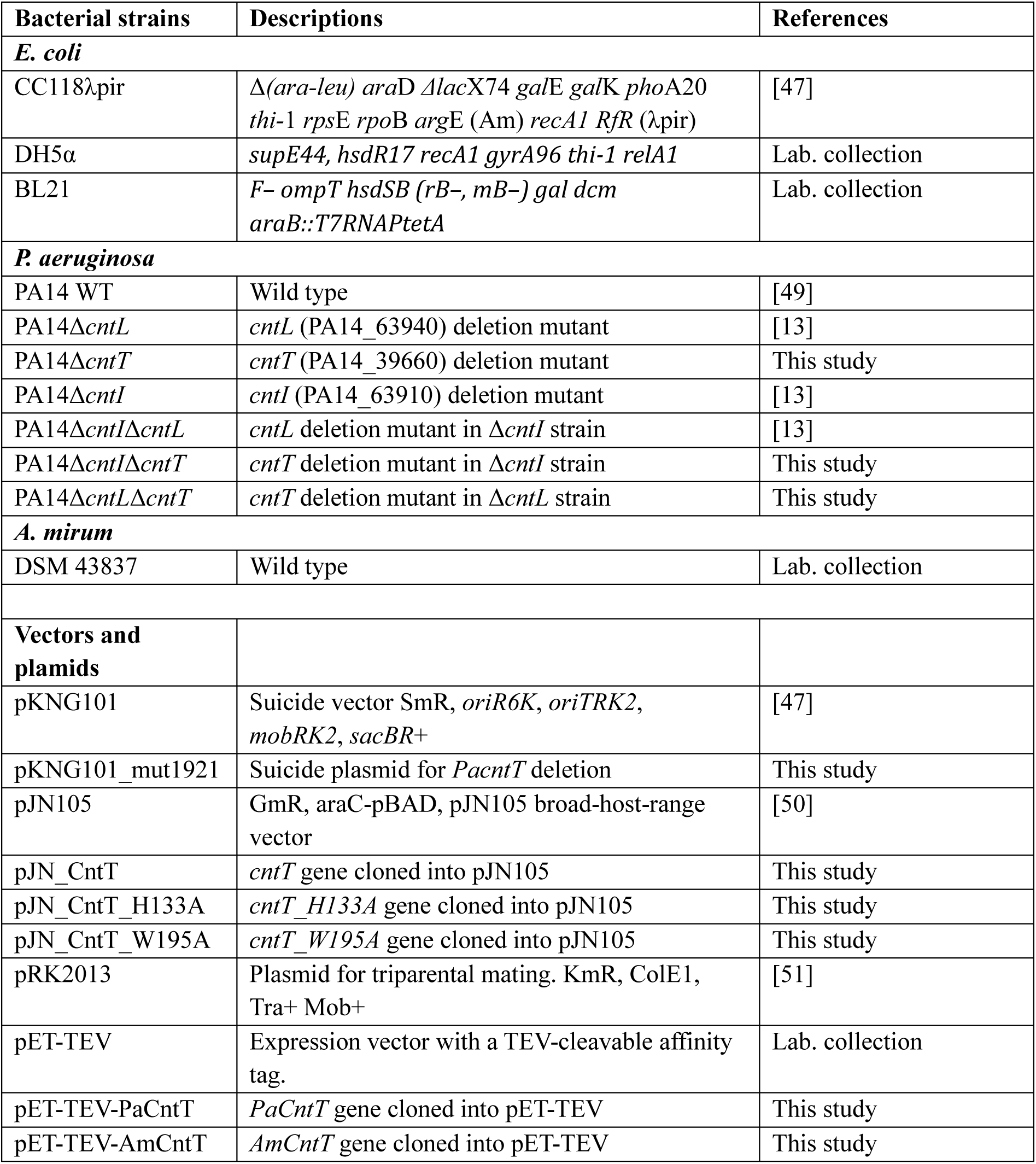
Bacterial strains, vectors and plasmids used in this study.

| Bacterial strains | Descriptions | References |
|---|---|---|
| ***E. coli*** |  |  |
| CC118λpir | Δ*(ara-leu) ara*D *Δlac*X74 *gal*E *gal*K *pho*A20 *thi*-1 *rps*E *rpo*B *arg*E (Am) *recA1 RfR* (λpir) | [47] |
| DH5α | *supE44, hsdR17 recA1 gyrA96 thi-1 relA1* | Lab. collection |
| BL21 | *F− ompT hsdSB (rB−, mB−) gal dcm araB::T7RNAPtetA* | Lab. collection |
| ***P. aeruginosa*** |  |  |
| PA14 WT | Wild type | [49] |
| PA14Δ*cntL* | *cntL* (PA14_63940) deletion mutant | [13] |
| PA14Δ*cntT* | *cntT* (PA14_39660) deletion mutant | This study |
| PA14Δ*cntI* | *cntI* (PA14_63910) deletion mutant | [13] |
| PA14Δ*cntI*Δ*cntL* | *cntL* deletion mutant in Δ*cntI* strain | [13] |
| PA14Δ*cntI*Δ*cntT* | *cntT* deletion mutant in Δ*cntI* strain | This study |
| PA14Δ*cntL*Δ*cntT* | *cntT* deletion mutant in Δ*cntL* strain | This study |
| ***A. mirum*** |  |  |
| DSM 43837 | Wild type | Lab. collection |
| **Vectors and plamids** |  |  |
| pKNG101 | Suicide vector SmR, *oriR6K*, *oriTRK2*, *mobRK2*, *sacBR*+ | [47] |
| pKNG101_mut1921 | Suicide plasmid for *PacntT* deletion | This study |
| pJN105 | GmR, araC-pBAD, pJN105 broad-host-range vector | [50] |
| pJN_CntT | *cntT* gene cloned into pJN105 | This study |
| pJN_CntT_H133A | *cntT_H133A* gene cloned into pJN105 | This study |
| pJN_CntT_W195A | *cntT_W195A* gene cloned into pJN105 | This study |
| pRK2013 | Plasmid for triparental mating. KmR, ColE1, Tra+ Mob+ | [51] |
| pET-TEV | Expression vector with a TEV-cleavable affinity tag. | Lab. collection |
| pET-TEV-PaCntT | *PaCntT* gene cloned into pET-TEV | This study |
| pET-TEV-AmCntT | *AmCntT* gene cloned into pET-TEV | This study |

**Table 3:**
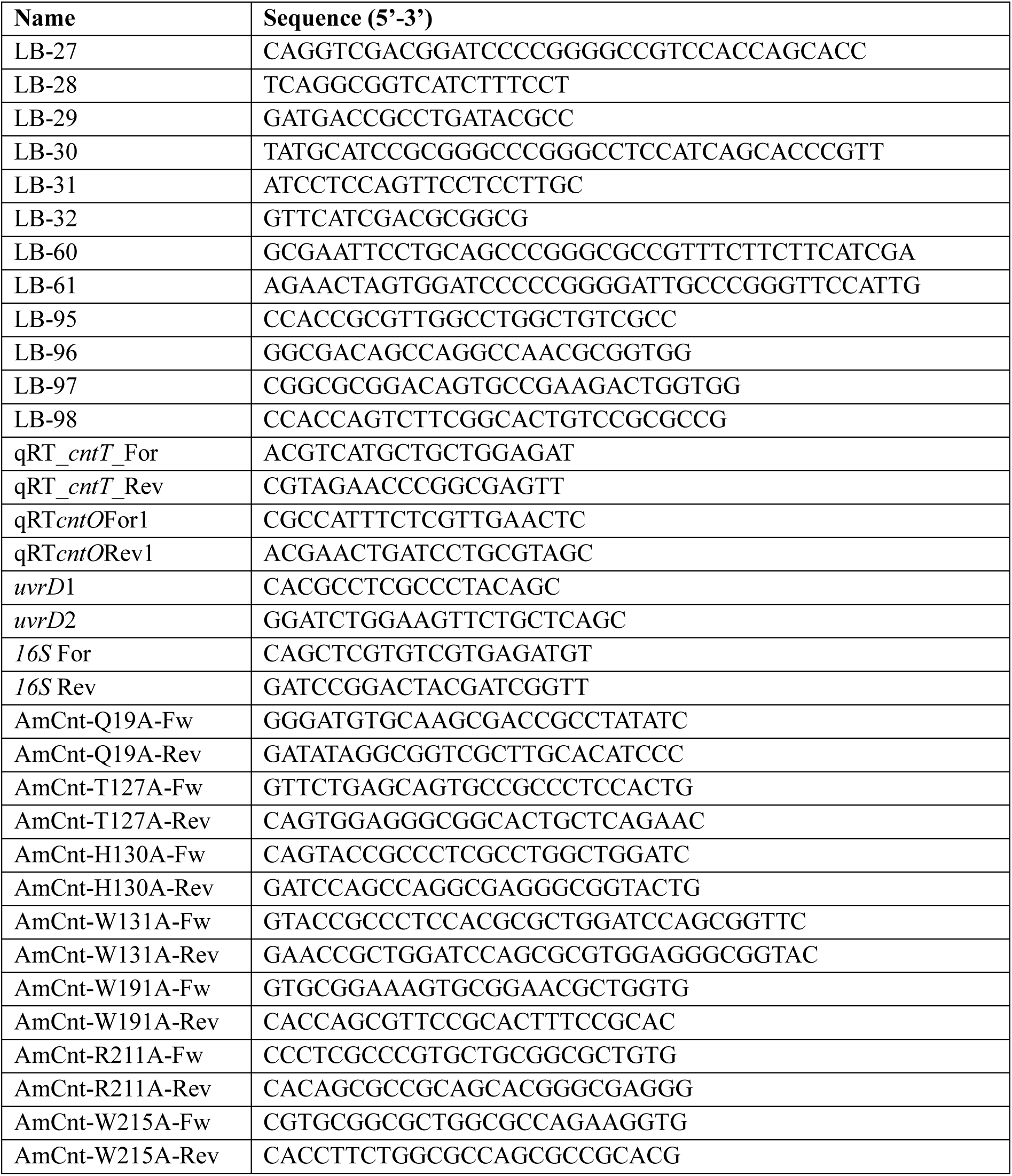
Primers used in this study.

### Site-directed mutagenesis of AmCntT

Site-directed mutagenesis was performed using the pET-TEV-AmCntT plasmid as a template. Seven single alanine substitutions were introduced at positions Q19, T127, H130, W131, W191, R211 and W215. Mutagenesis was carried out according to the manufacturer’s protocol for the QuikChange II Site-Directed Mutagenesis Kit (Agilent Technologies). The primer pairs used for each mutation are listed in Table 3.

### Statistics

Statistics were determined using one-way analysis of variance (ANOVA) followed by a Tukey’s post-test. Figures and statistics were performed using GraphPad Prism 10.4.1.

## ACKNOWLEDGEMENTS

We acknowledge Drs Yingjie Bai and Xiaoguang Lei (Department of Chemical Biology, College of Chemistry and Molecular Engineering, Peking University, Beijing, P. R. China) for chemical pseudopaline production and supply. We are gratefully to the whole RV group for constant support in the project. This work was supported by the grants: RF20210502871 from “Vaincre la Mucoviscidose” and “Gregory Lemarchal” association allocated to RV; ANR-20-CE11-0018 “Pseudo-Traffic” allocated to RV and PA and ANR-24-CE44-7195-04 “TonZyme” allocated to PA and RV.

## AUTHOR CONTRIBUTIONS

Conceive the experiments: LB, CL, LO, PA and RV. Perform, analyse and interpret the experiments: LB, CL, LO, TB, CB, GB, PA and RV. Write the paper: LB, CL, PA and RV. Secure funding’s: PA and RV.

## The authors declare no conflict of interest

## Supplementary Figures

**Figure S1:**
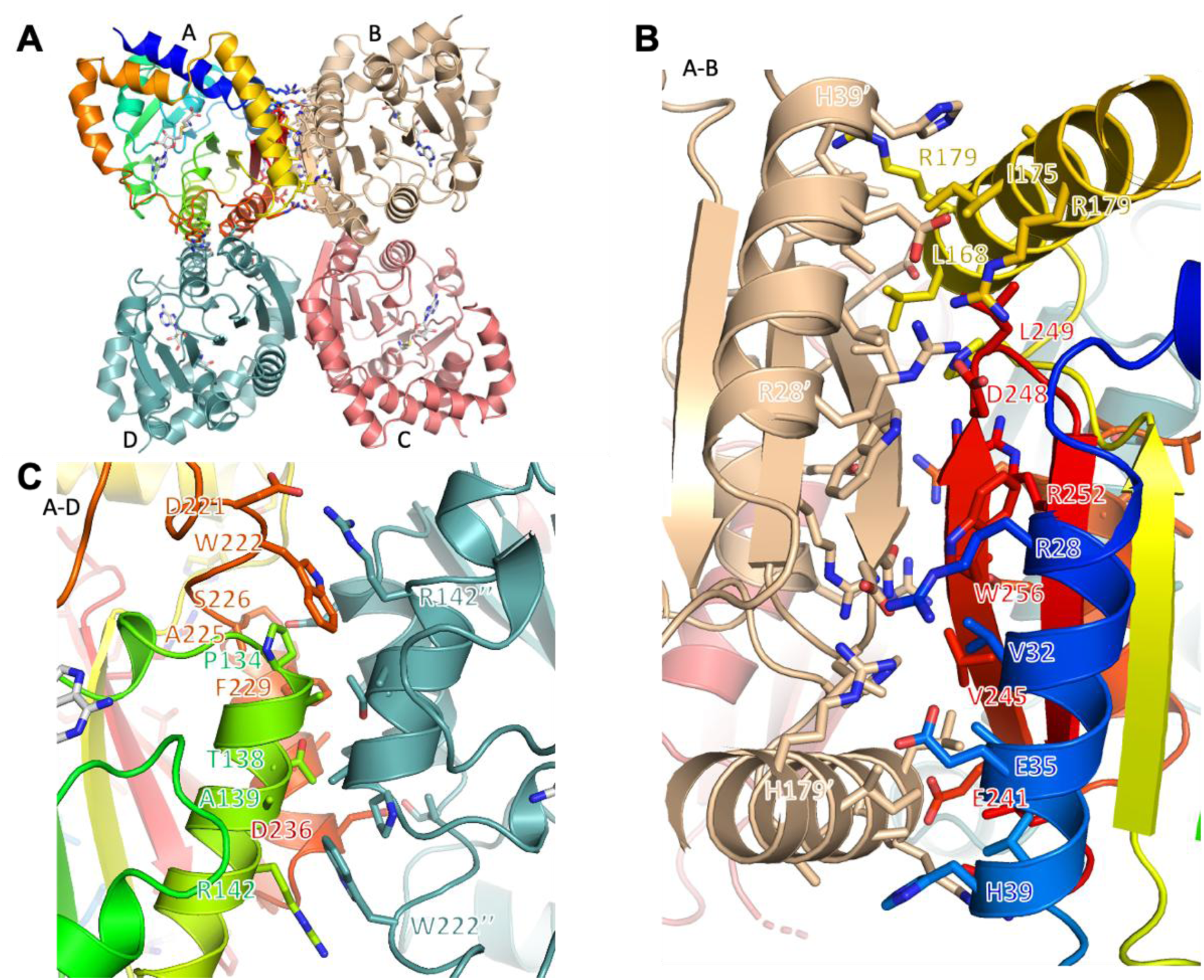
Assembly of the tetramer is mainly mediated by the last C-terminal secondary structure elements (α9β6β7). (**A**) The total buried surface area upon tetramer formation is 7997 Å2. (**B**) Details of the residues involved in the A-B interface. This interface represents 2886 Å2. (**C**) Details of the residues involved in the A-D interface. This interface represents 1214 Å2.

**Figure S2:**
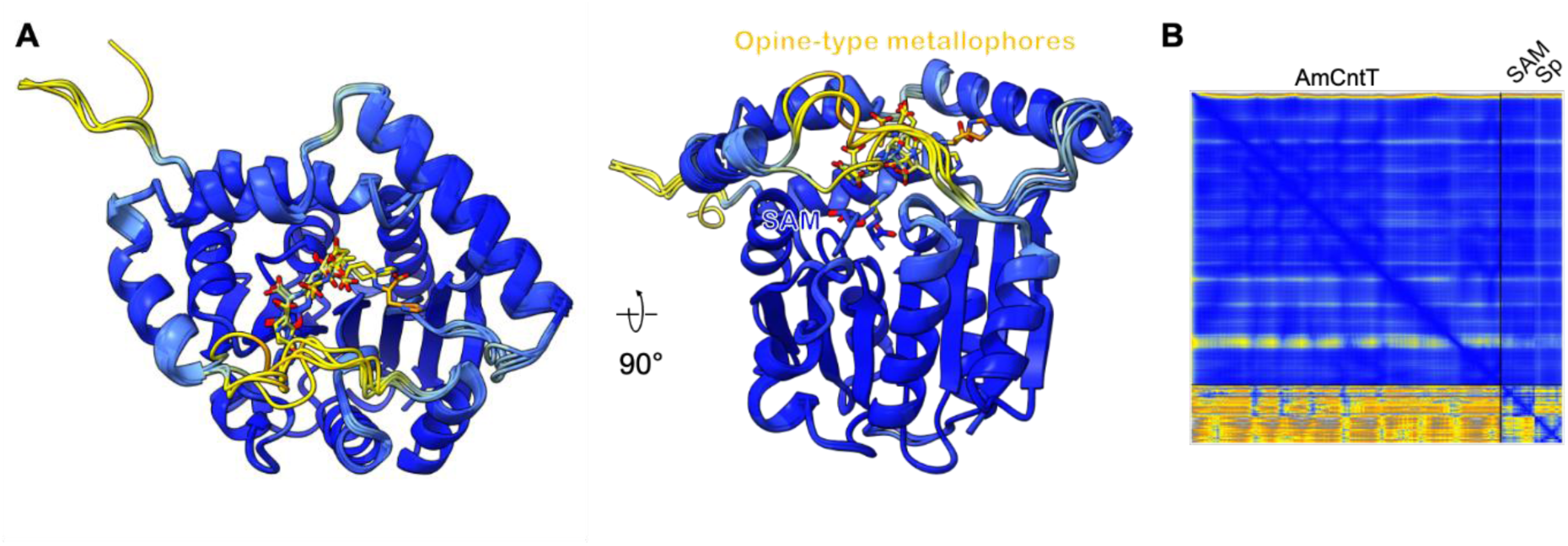
Boltz structure predictions of AmCntT in complex with SAM and the four opine-type metallophores. (**A**) Structure of the predicted complexes, showing that all four metallophores are predicted to bind in the same cavity, with a carboxylate moiety in proximity to the Sd atom of SAM. (B) AlphaFold Predicted Alignment Error (PAE) diagram for the AmCntT-SAM-Staphylopine (Sp) complex.

